# Incremental Self-Organization of Spatio-Temporal Spike Pattern Detection

**DOI:** 10.1101/2023.07.29.551088

**Authors:** Mohammad Dehghani-Habibabadi, Lenny Müller, Klaus Pawelzik

## Abstract

Nervous systems utilize temporally precise patterns of activity. However, the mechanisms by which spike patterns are processed are not known. In particular, the fact that during learning different patterns are distributed over time raises the question of how groups of neurons become selective for new spike patterns without overwriting already learned patterns. A simple one-layer spiking neural network model is presented that learns to recognize spatiotemporal spike patterns sequentially. The approach integrates biological synaptic mechanisms, including Hebbian learning, heterosynaptic plasticity, and synaptic scaling, allowing groups of neurons to self-organize selectivity for a set of spike patterns. Spoken words, transformed by a cochlear model into spatio-temporal spike patterns, are learned without supervision. This work suggests how the brain can use temporal spike codes and provides a novel, scalable, efficient, and noise-tolerant solution to the stability-plasticity dilemma.

## Introduction

In the brain, temporally ordered spike patterns may reflect temporal structures in external stimuli. First spike times also encode the amplitudes of their input spikes, giving rise to a rank order code^1^. Recently, it has been shown that objects and object categories are encoded in the sequences of spikes contained in short (∼100ms) population bursts whose timing is independent of the precise timing of external stimuli.^2^. Several more or less biologically plausible models have been proposed for reading spike sequence codes. For example, spike patterns could in principle be detected by adjusting appropriate transmission delays (^3, 4^), or with appropriate architectures and learning rules by recurrent spiking networks (^5, 6^). While these alternative approaches remain to be investigated, we focus here on the elementary finding that, with suitable synaptic efficacy, simple integrate and fire neurons are selective for spike patterns contained in short (∼50 - 100ms) epochs. This was first demonstrated by the introduction of the Tempotron^7^, which can discriminate between sets of patterns. However, the original Tempotron uses a supervisory signal which limits its biological plausibility. As an extension of the Tempotron, which does not provide any information about the timing of the output spike, Chronotrons have been discussed, which also use supervised learning algorithms to force neurons to fire at specific times during a pattern.^8–10^. Toward a more biological learning algorithm, as a network always receives input, it is required to be quiescent when receiving random input and to fire when a specific pattern is present, which is embedded in background activity. This can be achieved through correlation-based learning based on N-methyl-D-Aspartate (NMDA) receptors^11–13^ by selecting synapses with a sufficiently significant correlation between input and membrane potential^14^. The selection of synapses can also be done completely unsupervised based on pre and post-synaptic hetero-synaptic plasticity. In fact, it was already demonstrated in^15^that a combination of fundamental but realistic mechanisms, including synaptic scaling, Hebbian mechanisms, and hetero-synaptic plasticity, leads to a balance of excitatory and inhibitory inputs, and allows individual neurons to become detectors of repetitive patterns in the input without any supervision. There^15^ it was also found that the memory for the learned pattern is retained if the pattern is not shown again and only noise is presented afterwards which enables learning also of patterns repeating at low rates. If the learned embedded patterns are not presented to the neuron and the scaling term is large enough, all weights become scaled up until the neuron fires at the desired rate. This explains why memory is maintained when learning continues with random input patterns^15^. For the case of a single output neuron this approach, however, suffers a critical drawback: if a different embedded pattern is presented after learning a first pattern, this single neuron model encounters catastrophic forgetting: the memory for the original pattern becomes lost, and the new pattern is learned. A system with several output neurons has the potential to detect and identify multiple patterns^15^. However, it is not known by what mechanisms such a model could become selective by receiving the patterns to be learned one at a time. This amounts to finding a solution to the notorious stability-plasticity dilemma (SPD): Memory persistence depends on the stability of synaptic weight patterns in the neural networks that encode memories^16^. However, in order to learn from and adapt to new experiences, the brain must also be plastic. In fact, plasticity is a permanent mechanism and neurons constantly modify their synapses to fire at the desired biological firing rate^17, 18^. To understand learning, memory consolidation and retrieval, it is essential to understand how neural networks maintain stability in the face of ongoing plasticity. Deeper insights into the balance between stability and plasticity in biological neural networks also have the potential to lead to the development of more robust and efficient artificial neural networks. In particular, stable networks may fail to learn new information, while hyperplastic networks may suffer from forgetting, by replacing previously learned information with new information.^19–21^. The question therefore arises as to what mechanisms are required for a spike pattern detection neural network to be both stable and plastic in order to solve the SPD.^22^.

In this study, we use a modification of the model from^15^. First we show that also here a combination of realistic synaptic mechanisms enable a group of neurons to learn spatio-temporal spike patterns. Then we investigate the conditions for which the learned weight distributions remain particularly stable during ongoing plasticity when the learned patterns are absent in the input. Subsequently, we identify conditions for which new patterns are indeed learnable in assemblies of output neurons such that the weights related to previously learned patterns are maintained, which allows for incremental learning. Finally, we use spike patterns from spoken words to demonstrate that this model works also for real world data.

## Results

Neurons receive input from a large number of excitatory and inhibitory neurons. For simplicity, we use the example of 400 excitatory and 100 inhibitory input neurons which all fire with Poisson statistics and fixed rates of 5Hz and 20Hz, respectively. Embedded patterns are short epochs with the same statistics, however, contain consistent patterns which repeat either with frozen spike times or are noisy versions of a fixed pattern (Figure 1).

**Figure 1.**
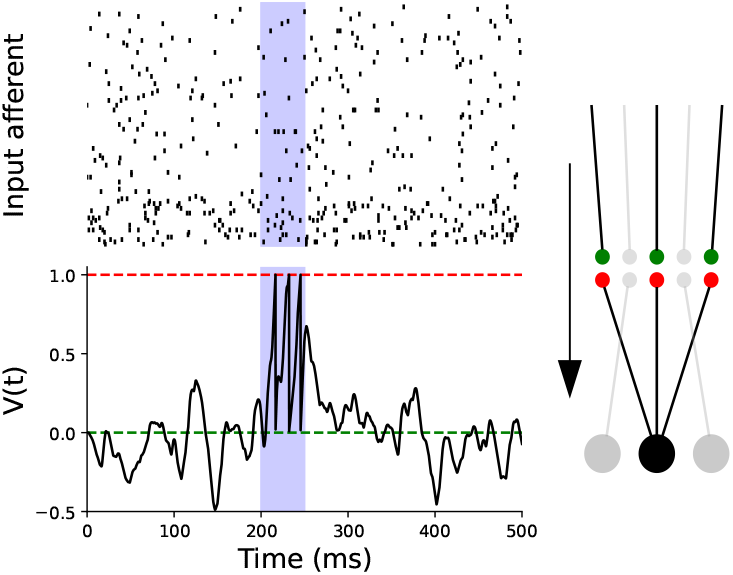
Learning of an Embedded Pattern. (Top) Input Activity: Spike trains from 500 afferent neurons, comprising 80% excitatory neurons firing at 5 Hz and 20% inhibitory neurons firing at 20 Hz. A 50 ms random pattern is embedded during the interval highlighted in blue. **(Bottom) Membrane Potential Response:** The black trace shows the membrane potential after learning, when the embedded pattern is present in the input. Learning enhances the neuron’s response during the embedded pattern interval (blue background) and globally balances fluctuations outside of the embedded pattern. As a result, the neuron generates spikes specifically during the embedded pattern period. The green line represents the resting potential, and the red line indicates the firing threshold.

The learning algorithm is taken from^15^ and sketched here for convenience (for details also see Materials and Methods). It combines plausible synaptic plasticity mechanisms for which the signs of the weights remain unchanged throughout the learning process, i.e. Dale’s law is enforced.

The algorithm incorporates homeostatic plasticity that is applied to excitatory synapses only and works to regulate the firing rate to match the biological firing rate^17, 18^. This means that when the firing rate of a neuron exceeds the desired rate on long time scales, all excitatory input synapses undergo a process of down-scaling, and they are scaled up when the firing rate is lower than desired. This synaptic scaling is crucial for the maintenance of system stability. However, because this mechanism is universally applied to all excitatory synapses, regardless of their efficacy, it is not sufficient to identify embedded patterns, that is, the neuron’s spike timing does not become pattern specific under synaptic scaling alone.

Hebbian mechanisms affect both excitatory and inhibitory synapses and depend on correlations between input kernels and deviations of the membrane potential from a given threshold. For changes in inhibitory synapses, weight enhancement is driven by positive deflections, while negative deflections contribute to their attenuation. For excitatory synapses, only positive deflections are allowed to contribute, mimicking NMDA-dependent processes. It is well understood that without any further constraints, this approach would lead to an instability of excitatory synaptic efficacy. To avoid this the efficacy of individual synapses is limited which is reasonable since there is a variety of biological constraints that prevent unlimited growth. Last not least also hetero-synaptic plasticity of excitatory synapses are taken into account on both, the pre- and the post-synaptic side where the strengthening of a synapse occurs at the cost of weakening others based on a signal of for synaptic increase (see Materials and Methods and^15^).

To determine whether a neuron has learned or memorized a particular pattern, the timing of the spikes in the post-synaptic neurons is considered. If all output spikes occur during the epoch when the embedded pattern is present in the input, one can conclude that the neuron has learned that pattern. To evaluate how well the learning mechanism and the memory recall work in the recognition of random patterns,the average percentage of spikes that correctly identify the patterns is determined, which is denoted by *R*,

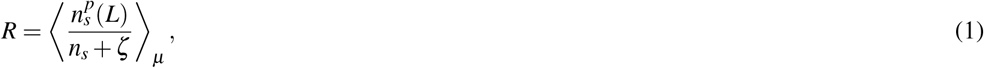

where *n*_*s*_ is the total number of spikes observed during a test period and 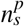 is the number of spikes related to the presence of the pattern to be detected. To account also for spikes that may occur shortly after the pattern due to the finite decay time of the excitatory synaptic kernel, the test window is extended by *L* ms after the pattern ends. To ensure a definitive result (*R* = 0) when no spikes are observed at all, a small arbitrary positive value *ζ <<* 1 is added to the denominator. The ratio is then averaged over an ensemble of simulations denoted by *µ*, where each input contains the embedded patterns and independent random background spikes.

### 1 Single Post-Synaptic Neuron

Here, we first reproduce the result from^15^ which showed that the mechanisms mentioned above enable a single post-synaptic neuron to learn embedded patterns. Since in this case only one postsynaptic neuron is taken into account presynaptic heterosynaptic plasticity does not participate (for details see Material and Methods). We find that also the present variant of the model in^15^ results in a balance of excitatory and inhibitory inputs into the target neuron which yields robustness of pattern detection against a range of stochastic perturbations. This is particularly significant when the training patterns are generated from Poisson processes with temporally modulated rates. Figure 1 shows the membrane potential of the neuron after learning a random fixed (frozen) pattern of length 50 ms. The embedded patterns used for the subsequent learning are taken from the same point processes for inhibitory and excitatory neurons, but are fixed and repeated in every stimulation. Figure 2A shows that the R-value, used as a measure of convergence, after learning reaches one (three spikes in the time window that the patterns was present and none outside). This shows that the neurons learn to fire only when presented with the embedded pattern and remain inactive otherwise. As in^15^ we find that the trained neuron retains its selectivity when after learning a specific pattern only random inputs are presented, and that this changes dramatically when instead of random spikes a different pattern is repeatedly presented: here the neuron looses its selectivity for the first pattern and becomes selective for the second.

**Figure 2.**
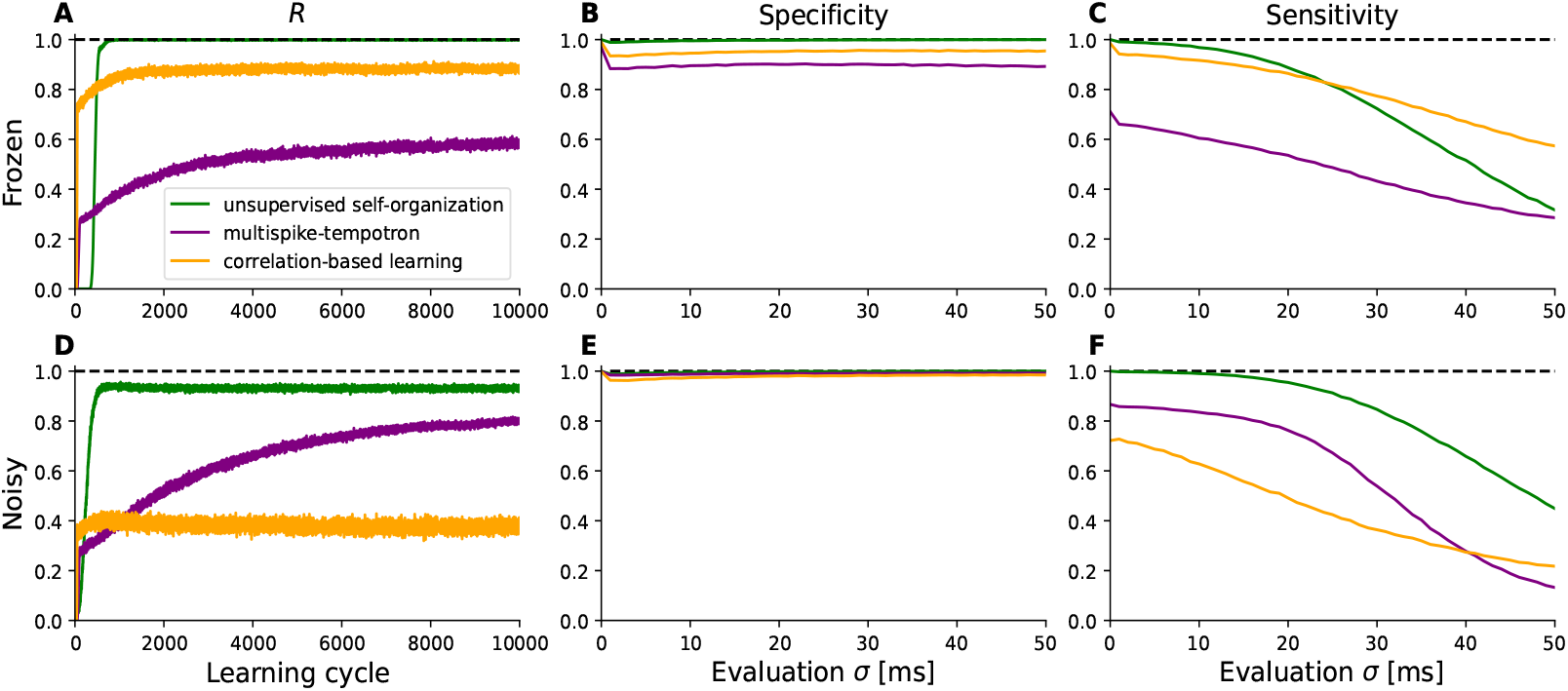
Comparison of Network Performance Using the *R* Metric. **(A) Training Performance with Fixed Input Patterns:** Performance of our unsupervised self-organizing model, pure correlation-based learning and the multispike-tempotron learning algorithm from a 2016 study on spiking neurons^14^ over 10,000 learning cycles using fixed (frozen) input spike patterns. The performance metric *R* represents the average percentage of postsynaptic spikes that occur during the embedded pattern interval, indicating the neuron’s ability to recognize and memorize the pattern without input variability. **(B) Specificity with Modulated Spike Patterns (After Training on Fixed Patterns):** Specificity of the networks trained in (A) when evaluated with temporally modulated Poisson spike patterns generated from the frozen patterns with varying temporal jitter *σ* . The modulated patterns are created by convolving each spike with a Gaussian distribution (mean zero, standard deviation *σ* ) to introduce temporal jitter, serving as the firing rate for a Poisson point process. As soon as temporal jitter is introduced the specificity of all models drops but other than that the specificity was unaffected by the strength of the temporal jitter in all models investigated. **(C) Sensitivity with Modulated Spike Patterns (After Training on Fixed Patterns):** Sensitivity of the networks trained in (A) when evaluated with temporally modulated Poisson spike patterns generated from the frozen patterns with varying temporal jitter *σ* . The strength of the temporal jitter strongly affects the performance of the all the models. Our model holds a near perfect sensitivity for small amounts of temporal jitter whereas for stronger jitter the drop in performance is much larger than in the compared models.**(D) Training Performance with Modulated Input Patterns (***σ* = 2 **ms):** Performance of our model over 10,000 learning cycles using input spike patterns modulated with a Gaussian convolution of standard deviation *σ* = 2 ms. This simulates learning under input variability resembling cortical codes, generating an ensemble of spatio-temporal patterns with Poisson statistics, including spike timing jitter, failures, and a Fano factor equal to 1. The performance metric *R* reflects the neuron’s ability to recognize and memorize the embedded patterns despite the input variability. Similar to (A) our model holds a superior performance over the compared models. The final *R*-value is below the one with frozen input patterns. Contrary to (A) the temporal jitter turned out to be helpful to multispike-tempotron learning and detrimental to pure correlation-based learning.**(E) Specificity with Modulated Spike Patterns (After Training on Modulated Patterns):** Performance of the networks trained in (B) when evaluated with modulated spike patterns using different temporal jitter widths *σ* . Slightly improved specificity for all models compared to the ones trained with frozen input patterns. **(E) Sensitivity with Modulated Spike Patterns (After Training on Modulated Patterns):** Performance of the networks trained in (B) when evaluated with modulated spike patterns using different temporal jitter widths *σ* . Sensitivity was improved in our model when training on noisy patterns. Close to optimal sensitivity for a broad range of temporal jitter widths. Results are based on an average of 500 simulations.

This model for synaptic plasticity is particularly robust with respect to a range of perturbations of the patterns, including jitter of spike times, failures and additional stochastic spikes (not shown). Most noteworthy is its performance for temporally modulated Poisson spike rates since they appear to represent cortical codes. Here it shows superior performance when compared with previous approaches as e.g. the learning algorithm described in a 2016 study on spiking neurons^14^ (Figure 2).

To generate a temporally modulated Poisson spike rate pattern from a basic frozen spike pattern, the given fixed time-coded spike input of each afferent was converted into rate-coded input by first convolving each spike with a Gaussian distribution characterized by a mean of zero and a standard deviation of *σ* representing the the temporal jitter. The resulting function then served as a modulated firing rate for a Poisson point process. This procedure generates for each basic pattern an ensemble of spatio-temporal patterns with the statistics of a Poisson point process including failures and a Fano factor equal to 1. Patterns from this ensemble were then used for plasticity and determination of the robustness of pattern detection.

In Figure 2 the left column shows the performance during the learning process and the center and right column present an evaluation of the performance after learning for a range of different jitter widths of the modulated spike rate patterns.

We compared our model to two algorithms from^14^, pure correlation-based learning and multispike-tempotron learning. In contrast to our approach those models employ also a momentum heuristic and furthermore some attenuation^14^ in addition to utilizing a supervised learning rule. Even with these features these alternative models fail to reach the same performance level as the here introduced unsupervised self-organization for the case of frozen input spikes (i.e. no blurred Poisson inputs). We evaluated the network with modulated input spike trains (Figure 2B&C). Besides an overall drop immediately after *σ* = 0 the specificity was only little affected by the jitter width in the examined range in all models. The sensitivity was more affected where our approach held a better performance for smaller values of *σ* . The stronger decline of sensitivity for higher *σ* compared to the more linear decline in performance of the compared models is a hint that our model better incorporates the temporal structure of the patterns. When exposed to modulated spike rate patterns, the neuron takes longer to learn the embedded pattern, and the performance becomes less distinct, while still remaining close to *R* = 1. Compared to the other models, however, learning is faster. When exposed to higher levels of temporal blur of the patterns than present during learning, our model allows for a much greater noise robustness of the neuron (see Figure 2E&F). The reason for this stable performance lies in the balance between excitatory and inhibitory inputs which leads to more sparsely distributed weights (maintaining membrane potential fluctuations around the resting state, *V*_0_ (Figure 1)). This then enforces a stronger robustness against additional spikes fired or transmission failures which are mimicked by the missing of input spikes in the Poisson process. This model also holds a robust and stable memory trace for single patterns but suffers catastrophic forgetting when instead of the learned pattern a different pattern is embedded into the input^15^. For this several output neurons are needed which will be discussed in the following section.

### 2 Neural Network

In principle, a single neuron is capable of learning more than one embedded pattern along the lines of the Tempotron^7^. However, a system with only one post-synaptic neuron cannot identify when and which pattern is present in the input. This would require an ensemble of output neurons which acquire different selectivities. It was already shown that for this purpose pre-synaptic hetero-synaptic plasticity is essential^15^ since it induces a competition among out-going synapses. For investigation of the conditions under which plasticity and stability of memories meet, we consider a very simple network with several post-synaptic neurons and include also pre-synaptic hetero-synaptic plasticity.

We find that including this establishes approximately orthogonal weight vectors. As a result, when a new pattern is introduced, it adds an additional dimension to the system. The synaptic competition induced by pre-synaptic hetero-synaptic plasticity is sufficient to faithfully represent pattern identity and order in an ensemble of target neurons with shared input from a set of input neurons if, during learning, all patterns are presented in each epoch. To quantify this, we use a matrix-based approach to evaluate the performance of the population of neurons. The matrix has rows representing neurons and columns representing patterns, with matrix elements set to 1 if a neuron is active for a particular pattern and 0 otherwise. The rank of this matrix reflects the number of linearly independent row vectors. We propose a performance measure, denoted Ω, which is the ratio of the matrix rank to the number of patterns. When the rank of the matrix is equal to the number of patterns, it indicates that the population of neurons accurately represents the presence and order of the patterns in a stimulus. In this study, Ω is utilized as a criterion for evaluating learning, plasticity, and stability. As a simple example, we trained a network with seven post-synaptic neurons using four embedded patterns in each training epoch. We found that when the pre-synaptic competition was present, all four patterns were selectively represented by the neurons, regardless of whether the patterns were shown in every learning cycle or only stochastically (Figure 3). In the absence of pre-synaptic competition, the selectivity for the patterns was not effectively distributed among the neurons^15^. Therefore, hetero-synaptic plasticity promotes the functional specialization of output neurons to different subsets of patterns, facilitating the self-organization of a precise representation of the “which” and “when” aspects of input patterns by the neuronal ensemble.

**Figure 3.**
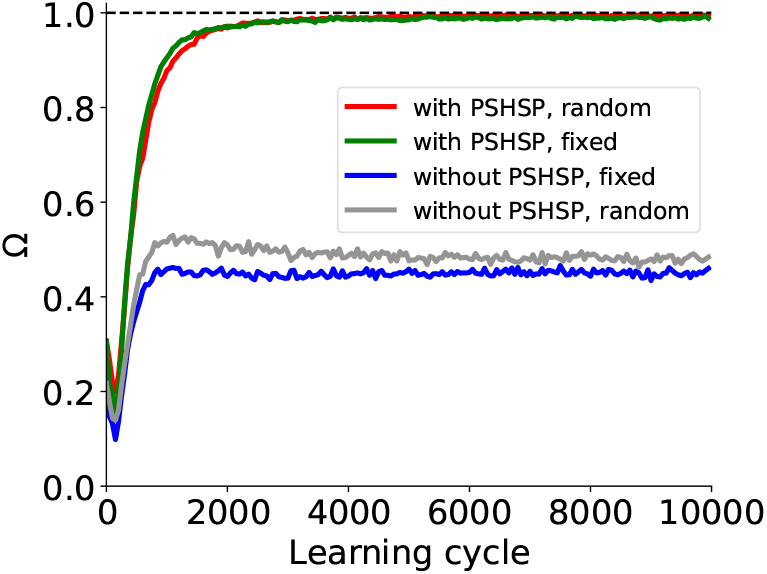
Representation of four embedded patterns by seven post-synaptic neurons. Ω versus learning cycle (There are 7 post-synaptic neurons). Green and Blue lines: Each pattern is shown in each learning cycle in a fixed position. Red and Gray lines: Each pattern is shown in the epoch with the probability of 1*/*2 at a given position. Note that in this case some learning cycles have no embedded pattern in the epoch. (The data are from every 50’th learning cycles. This figure is based on the average of 500 simulations.)

The fact that specialization (i.e. Ω → 1) takes place also when each pattern is shown intermittently (i.e., not in each epoch, the red line in Figure 3) indicates the robustness of memories not only against the absence of patterns but also against the presence of different patterns. This raises the question under which conditions true incremental learning is possible without overwriting previous memories. We find that this is indeed the case, however only when the patterns are given as Poisson rate modulations (as explained above), as opposed to precise pattern repetitions.

In Figure 4, the initial learning phase involves *M* = 7 target neurons learning two embedded patterns. After 30,000 learning cycles a third pattern is embedded but the previous patterns are absent in the input. The system’s ability to recognize all patterns is tested every 50 cycles (with plasticity switched off) to assess memory retention. Subsequently after another 30,000 learning cycles a fourth pattern is shown, again with the previous pattern not being presented anymore during learning.

**Figure 4.**
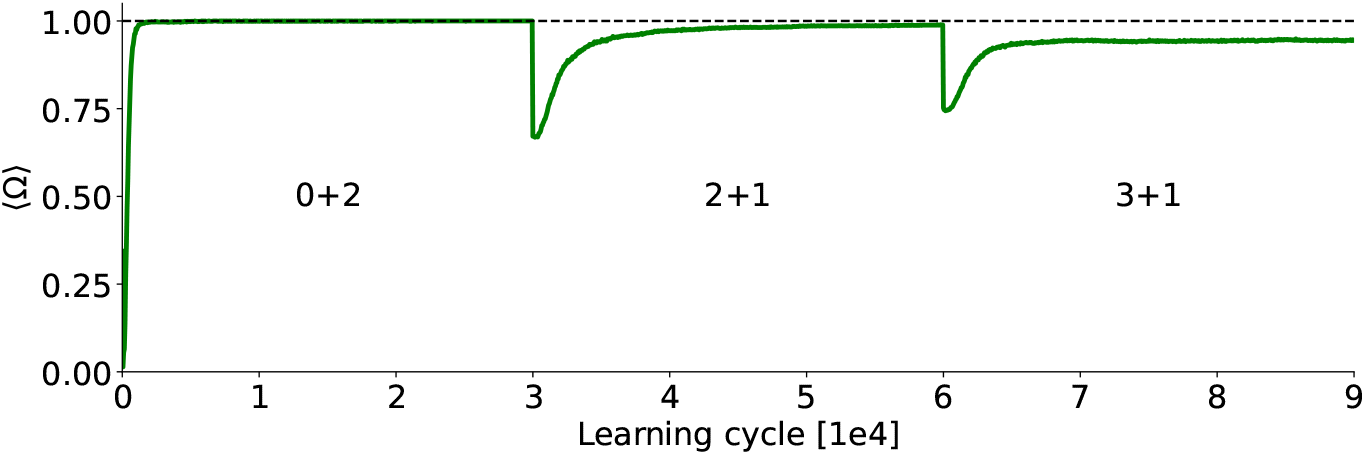
Learning curve in the incremental learning task. The network was initialized with 2,000 learning cycles of random input. In the first stage (0+2) there are two embedded patterns in the input for 30,000 learning cycles. After that the network is presented with a third pattern (2+1) but the first two patterns are no longer presented. After another 30,000 learning cycles a fourth pattern is embdded (3+1) but the third pattern (as well as the first two) are absent in the input. Every 50 learning cycles learning is stopped and Ω is computed for all the so far embedded patterns. There are 7 post-synaptic neurons and the results are averages over 500 simulations.

The drop at the 30,000 and 60,000 learning cycle marks is due to the new input pattern being incorporated which the network was just presented to and therefore did not yet learn. The increase back to Ω = 1 in the second stage (2+1) therefore resembles the learning of the third pattern while also demonstrating that the distributed selectivity for the first two patterns is maintained. To understand the superior stability of memories for patterns when learned from temporally modulated Poisson rate patterns we investigated the weight matrix *w* = *ab* with its pre- and postsynaptic components *a* and *b* respectively (see Materials and Methods). In particular, we tested the hypothesis that stability relates to the orthogonality of the weight vectors. As a graded measure *O* of ‘orthogonality’ we consider the volume spanned by the *M* weight vectors in the *N*^*E*^ -dimensional input space (*N*^*E*^ : number of excitatory afferents). We normalized the weight vectors leading to a matrix *Y* . The volume is then given by the product of the square roots of the *M* largest eigenvalues *λ*_*i*_ of *X* = *YY*^*T*^ :

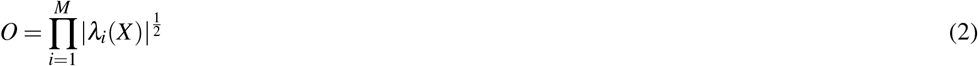

If the *M* normalized vectors are all mutually orthogonal this volume is *O* = 1, if one or more weight vectors are linearly dependent on the others this volume collapses to zero (*O* = 0). Figure 5A shows that for the case of jittered noisy (Poisson rate) patterns *O* is significantly increased compared to the case when two rigid patterns are shown. This difference allows the network to store the information about the patterns more effectively and leads to higher capacity of learnable patterns. When there are only two or three patterns present the network is able to learn to identify them regardless of frozen or noisy input. But with increasing load there is less space which leads to a point where network trained on frozen input patterns can not store all the information necessary (see Figure 5B). To demonstrate the scalability of the network we show the averaged Ω (Figure 5C) at the end of each incremental learning step for a range of postsynaptic neurons. We find that the a higher number of postsynaptic neuron indeed increases the performance in the respective learning steps altough not in a linear manner whereas the overall performance decay with the increasing number of presented patterns seems to be linear. To illustrate how the systems simultaneously maintains stability and plasticity we looked at the development of the weight vectors throughout the incremental learning task. We evaluated the cosine similarity of the weight vectors of each output neuron with its respective final state as shown in Figure 6 for an exemplary simulation. In each stage (every 30,000 learning cycles) there are vectors strongly shifting, indicating that their selectivity is changing. What is notable is that once a neuron has developed the same selectivity as in the final state, it is not pushed out of that regime anymore indicating that selectivity is stable.

**Figure 5.**
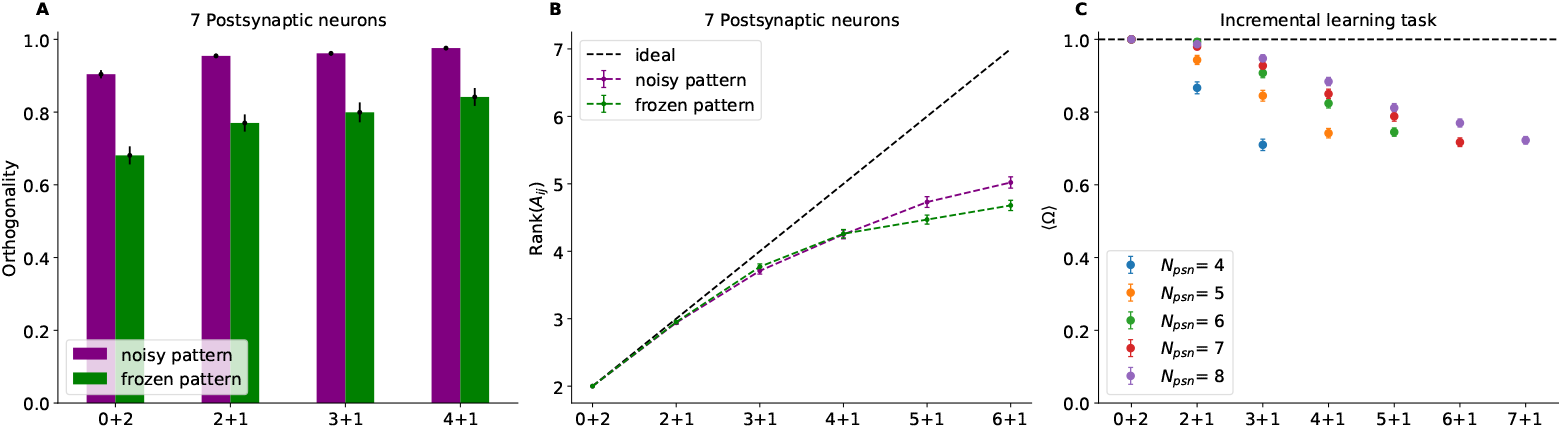
**A:** Orthogonality for original fixed patterns and Poisson spike rates. **B:** Ω **in the incremental learning task for differnt numbers of postsynaptic neurons. (A)** Purple: Poisson spike rates. Green: Initial frozen pattern. The computation of the orthogonality is performed with respect to the combined synpatic efficacy, *w* = *a*·*b*. There are 7 post-synaptic neurons. The results are averages over 100 independent simulations. **(B)** Rank of the activity matrix at the end of each incremental learning step for the same simulations as in A. **(C)** Performance in the incremental learning task with standard error. Ω is comptuted in every stage for all the patterns shown up to the respective stage. The results are averaged over 100 independent simulations.

**Figure 6.**
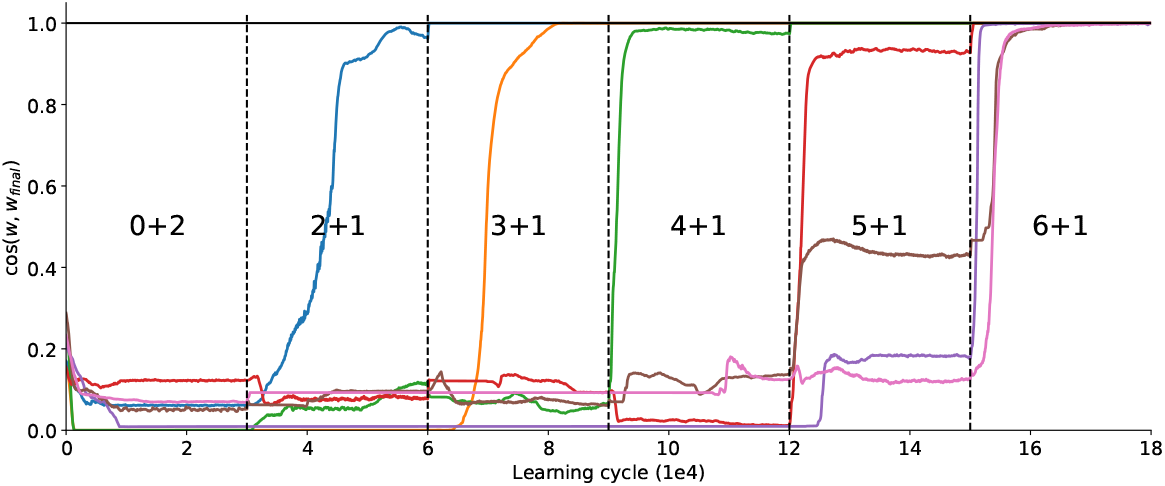
Cosine similarity between the weight vectors of the seven output neurons of an exemplary simulation during learning and their respective final state during incremental learning. The neurons develop selectivities at different points during learning and once they have, they remain fixed until the end of the evaluated learning cycles. Every 30,000 learning cycles a new pattern is shown while the previous patterns are not shown again.

As in^15^, we apply hetero-synaptic plasticity and synaptic scaling simultaneously. Introducing noise during the learning process can sometimes cause important spikes to be missed or generate additional spikes in afferent neurons. The missing of spikes primarily contributes to memory retention of previously learned patterns. When critical spikes are occasionally missed due to noise, the input-output correlation across synapses becomes more uniform. This uniformity enhances the effect of synaptic scaling over hetero-synaptic plasticity, allowing the synaptic weights associated with previously learned patterns to be preserved, thereby maintaining memory stability.

Conversely, the addition of spikes, especially in afferent neurons that are more correlated with the post-synaptic neuron’s membrane potential, facilitates the learning of new patterns. The increased activity in these highly correlated afferent neurons leads to selective strengthening of their synaptic weights through hetero-synaptic plasticity. This selective reinforcement enables the post-synaptic neuron to adapt effectively to new information, enhancing the network’s plasticity.

This mechanism demonstrates that while hetero-synaptic plasticity is essential for learning new patterns by reinforcing relevant synaptic connections, synaptic scaling—amplified by the occasional missing of spikes due to noise—is crucial for retaining previously learned patterns in the former model^15^ (not shown) and helpful in the model used here. The simultaneous application of both mechanisms allows the network to balance stability and plasticity, effectively incorporating new patterns without erasing existing memories. These findings provide valuable insights into how neural systems can preserve past information while remaining adaptable to new inputs.

### Neural Decoding of Spiked Audio

The aim of this section is to assess the algorithm’s ability to identify embedded patterns within signals that replicate natural conditions. We used auditory data from the Audio Signal dataset, which consists of spoken digits from 0 to 9 in English, converted into spike patterns by the Lauscher artificial cochlea model^23^. This method allows us to analyze spikes across 700 input channels, taking advantage of the high-quality, aligned studio recordings in the Spiking Heidelberg Digits (HD, Spiking Heidelberg Digits: SHD) dataset. The data set was selected for its suitability for audio-based classification. Moreover, this dataset includes recordings from 12 different speakers, providing a diverse basis for testing the model’s pattern recognition capabilities under naturalistic conditions.^23^

Figure 7 shows the amplitude and spatio-temporal spike patterns of speaker number 2 when saying “*four*” (panels A and C) and “*five*” (panels B and D), as represented in the files *lang-english_speaker-02_trial-6_digit-4* and *lang-english_speaker-02_trial-31_digit-5*, respectively.^23^

**Figure 7.**
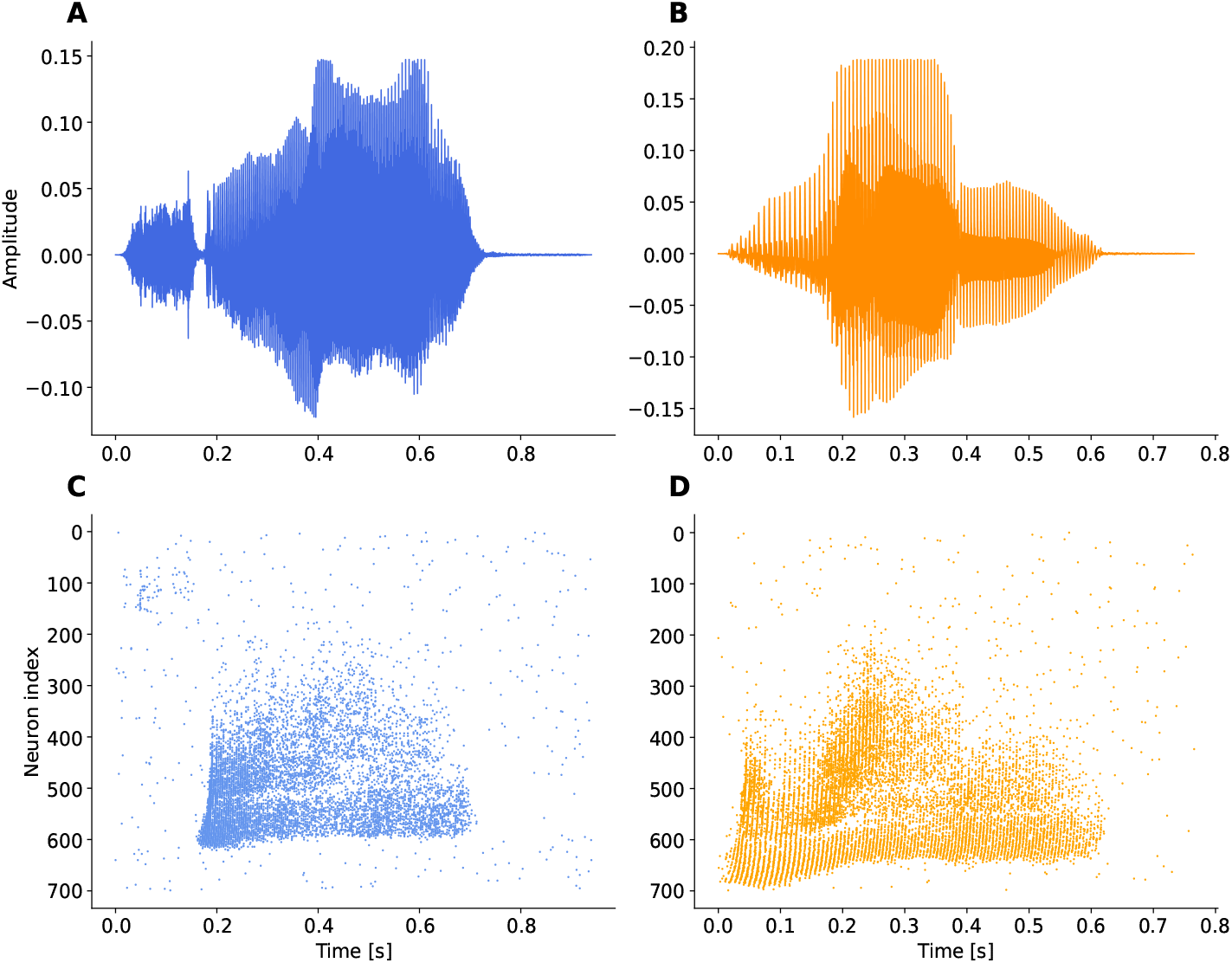
Spatio-temporal data in spike pattern of speech data. In Panel A, Speaker 3 says”*Zero*” (violet) and “*five*” (orange) in Panel B. Panels C and D show the spatiotemporal spike patterns corresponding to A and B, respectively. lang-english_speaker-02_trial-6_digit-4 and lang-english_speaker-02_trial-31_digit-5.^23^

These patterns, however, exhibit a high degree of synchronization, which poses the challenge of generating the same level of correlation between input and output across different afferents, potentially leading to identical synaptic efficacies. Therefore, the competition among excitatory neurons may need to be more sensitive to noise and small differences between afferents. To address this issue, we have improved the effectiveness of presynaptic heterosynaptic plasticity mechanisms (Materials and Methods). This adjustment enables a more subtle and effective modulation of synaptic strength, which is essential for discriminating between highly synchronized inputs.

Furthermore, these data are challenging because a single word can contain multiple phonemes, increasing the possibility that the network interprets each segment as a separate pattern and does not learn another word at all. Consequently, the network may allocate all spikes to a single stimulus, thereby overshadowing the subtle differences between phonemes. We trained networks with 6 post-synaptic neurons for five different word pairs ([“*Zero One*”], [“*Two Three*”], [“*Four Five*”], [“*Six Seven*”], [“*Eight Nine*”]) on two different conditions. For the first one from Fig. 9 we selected 10 trials from each of the 12 speakers for one word and combined them randomly with the other word from one of 10 trails from the 12 speakers. This training set of 120 pairs contained pairs with trials from different speakers as well as pairs where both words were trials from the same speaker. These 120 word pairs were shown 50 times with plasticity after every word pair so a total of 6,000 learning cycles.

**Figure 8.**
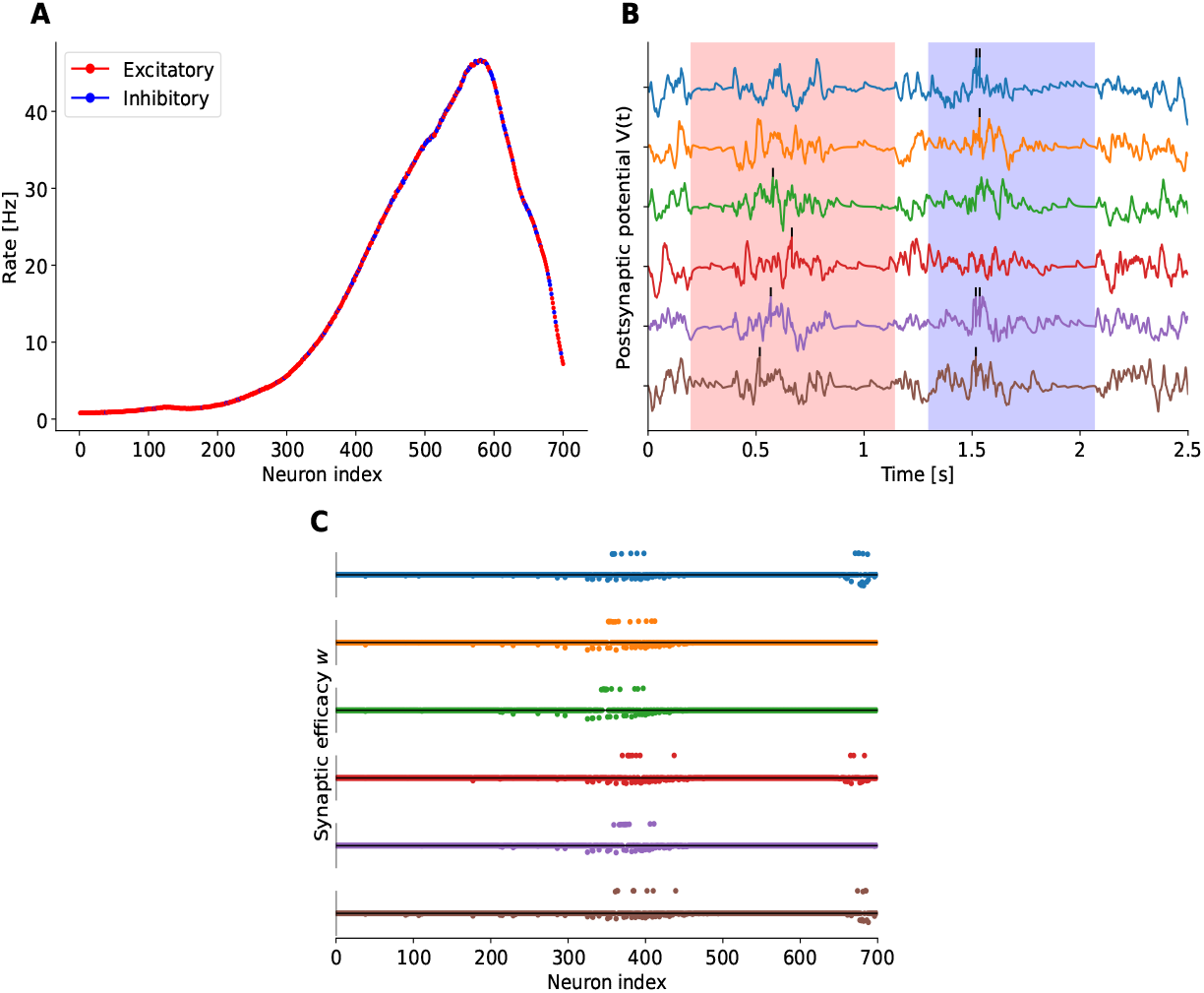
Application to speech data. Each embedded pattern was placed between background noise generated from the distribution in panel A (blue is inhibitory and red is excitatory). The network was trained to recognize only the spike patterns for “*zero*” (red background) and “*one*” (blue background) from speaker 3 and 4, trial number 4 and 9 respectively. B shows the membrane potential of each neuron after 6,000 learning cycles and the corresponding weight vector in panel C. The epoch length is 2500 ms, and the desired number of spikes is 2.

**Figure 9.**
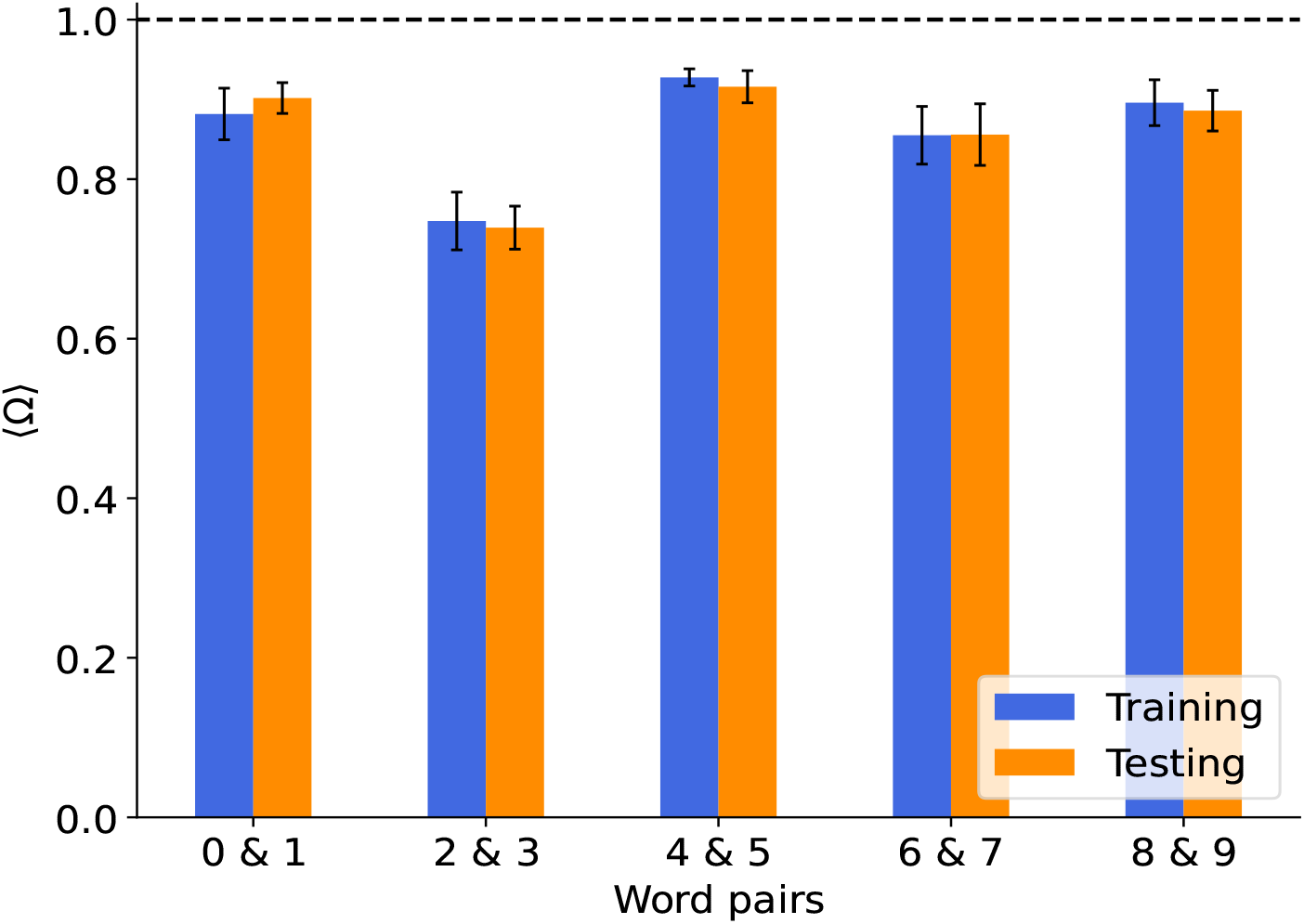
Training with trials from all speakers. The network is trained with a given word pair containing trials from randomly chosen speakers. The test set consisted of trials different form the training set. Each pair of words is presented to the network for a total of 6,000 learning cycles. The desired number of spikes is 2, and the number of postsynaptic neurons is 6. The results show the mean performance of 10 runs with different random combinations of the speaker trials in the training set.

After training the networks were evaluated on a test set which also consisted of 120 word pairs from random speakers but the individual trials were explicitly different ones that used in training. The similarity between training and testing performance in Fig. 9 shows that not only learning of the words occurred but also a generalization that allowed for successful identification of the other trials. Note that before embedding patterns, the network undergoes a preparation phase of 500 cycles of noise exposure, during which the desired firing rate of 0.8 Hz was established. For illustration of a trained network of 6 neurons, we show the membrane potentials when speaker number 3 said “*Zero*” and speaker number 4 “*One*” (Figure 8B). The corresponding weight vectors of this model (Figure 8C) reveal that distinct afferents contribute to the generation of spikes (black lines in Figure 8 B).

To further assess the generalization we trained networks on the second condition where all the trials used for learning came from one speaker. Here 20 trials from a chosen speaker were shown 300 times to again train for 6,000 learning cycles. After training a test set of 20 trails from each of the unused speakers was presented to the network. The averages over all the networks for the word pairs are shown in Fig. 10. here the standard deviations indicate that in this training condition the speaker used for training had a large influence on the performance of the network. In both training conditions differences in the word pairs were noteable ([“*Two Three*”] showing the worst performance in both conitions for example). This may be due to the allocation of spikes to the same phonemes of these are the most dominant features in both words as already mentioned. In our tests we found that two words can be learned also one after the other indicating that the incremental learning works also for other realistic data.

**Figure 10.**
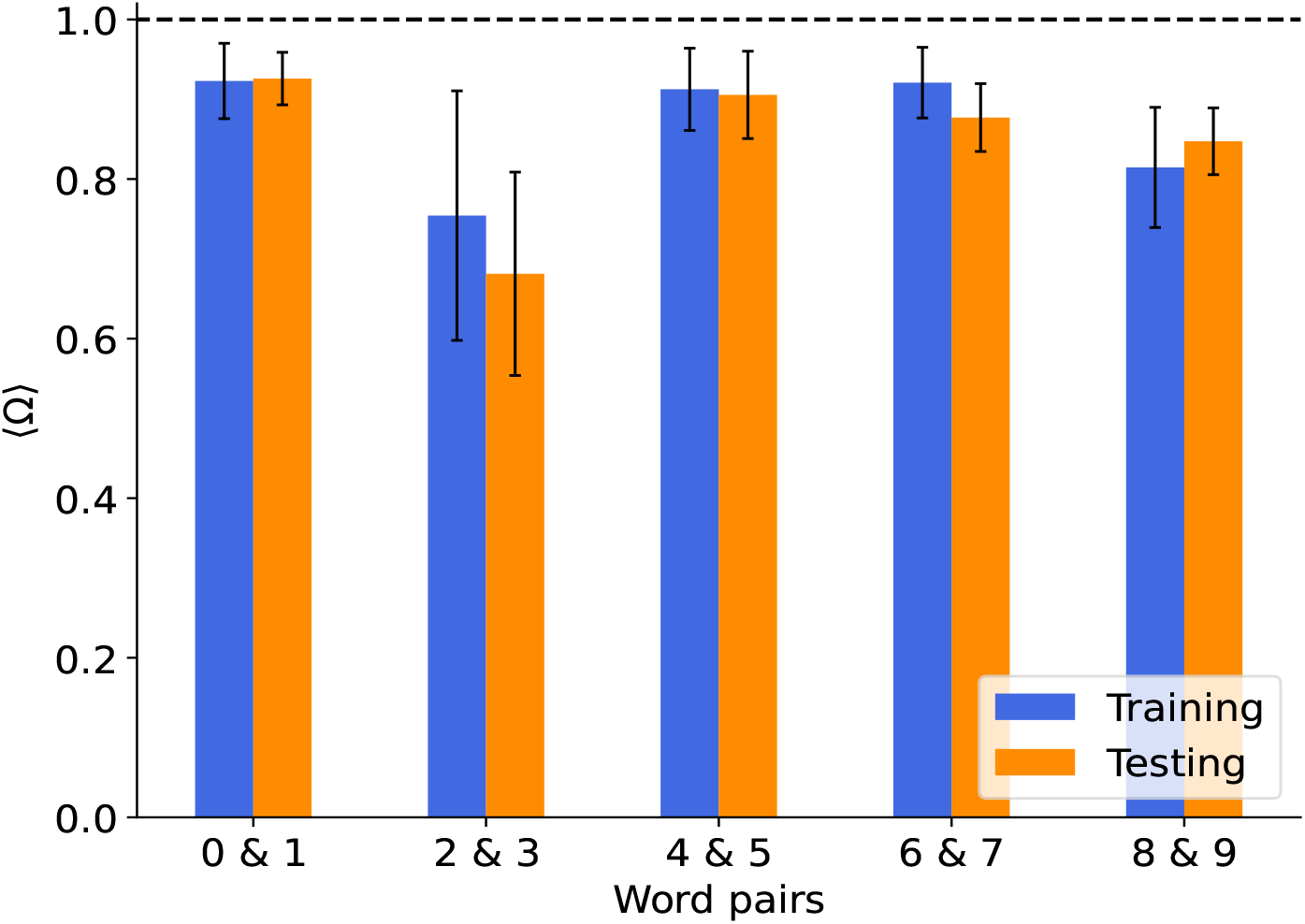
Training with trials from a single speaker. A set of trials from a single speaker was presented during training. The testing set consisted of trials from all other unused speakers. Each pair of words is presented to the network for a total of 6,000 learning cycles. The desired number of spikes is 2, and the number of postsynaptic neurons is 6. The results show the mean performance of 12 runs, trained on one of the 12 speakers, respectively.

## Discussion

Selectivity for spatio-temporal patterns can self-organize in simple one-layer networks of integrate and fire neurons^24, 25^. Particularly, a combination of Hebbian plasticity mechanisms, synaptic scaling for excitatory synapses and hetero-synaptic plasticity was demonstrated sufficient for robust spike pattern detection^15^. By using a variant of this model we found that the competition for weight increases of the pre-synaptic components of excitatory synapses which are controlled by the pre-synaptic neuron (termed ‘pre-synaptic hetero-synaptic plasticity’) have a tendency to orthogonalize the weight vectors of the target neurons in input space (Figures 3, 5A). When the patterns are supposed to be learned one after the other, our simulations demonstrate that the weights of the output neurons of a network stay more orthogonal if the patterns are stochastic variants of the original pattern. We showed this by generating the training patterns from a Poisson process with temporally modulated firing rates obtained from convolving the originally fixed pattern with a Gaussian function. In this case, orthogonality of the weights is higher also when the patterns are learned incrementally. The orthogonality then supports minimal interference with previously learned weight vectors^26, 27^.

Taken together, we proposed a biologically realistic solution to the stability-plasticity dilemma for networks that learn to detect spatio-temporal spike patterns without any supervision. This approach relies on the balance between excitatory and inhibitory inputs that naturally emerges during learning^15^ which was suggested to be necessary for maintaining memory already before^28^. To our knowledge this is the first approach towards a biologically realistic model for incremental self-supervised learning of spatio-temporal spike patterns. Previous approaches were proposed for recurrent networks^29, 30^ and rate-based models^26^ which relied also on systematic orthogonalization of weight vectors, however by different mechanisms. Furthermore, our approach relies heavily on mechanisms for synaptic scaling which has also been proposed before as a mechanisms for stabilizing long term memories, however in rate based models and with a *w*^2^ dependence of scaling^30^ which in our case was found to be detrimental for performance of learning (not shown).

Last but not least, our results align with previous research^31^ emphasizing noise in the input for supporting incremental learning by driving weights to become more orthogonal and serving to minimize their overlap for different output neurons. When tested on auditory data the approach was found to get along with the variability of real data suggesting that a one layer network that is trained incrementally with unlabeled data could be sufficient for speech recognition.

## Materials and Methods

The one-layer network consists of *N* pre-syaptic input neurons and *M* post-synaptic output neurons. The membrane potential of the output neuron *j* is computed from the external input *I*_*j*_(*t*) using the leaky integrate and fire equation

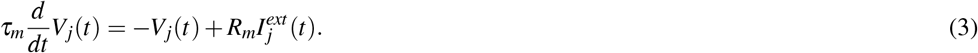

The external signal is given by the spikes from the input neurons with an alpha-function shape

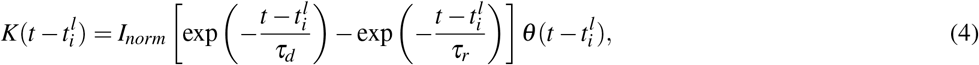

where *τ*_*r*_ and *τ_d_* denote the rise and decay time constant respectively. The term 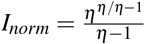, where 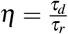 , normalizes the amplitude of the kernel *K* to unity. The input current is then given be the sum over all input neurons of the summed spikes of an individual neuron multiplied with the respective weight *w*_*ji*_ of that neuron

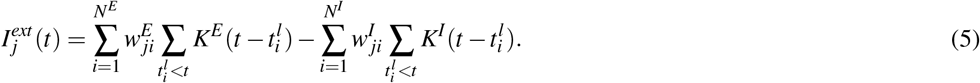

Note that the superscript denotes whether it is an inhibitory neuron (I) or an excitatory neuron (E). Signal transmission is more effective when pre-synaptic neurons release more neurotransmitters and fire more frequently, and also when post-synaptic neurons have more receptors and sensitivity. Therefore, the efficacy of a synapse,*w*_*ji*_ (pre-synaptic neuron *i* targeting a post-synaptic neuron *j*), can be expressed in terms of the combination of pre-synaptic and post-synaptic components

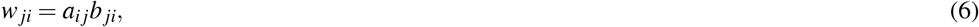

where the resources for *a*_*ij*_ are provided by the pre-synaptic neuron *i* and *b* _*ji*_ is supported by the post-synaptic neuron *j*.

The efficacy of a synapse can be modified separately in the pre- and the post-synaptic components through synaptic plasticity, resulting in changes in the total efficacy of the synapse in transmitting signals between neurons.

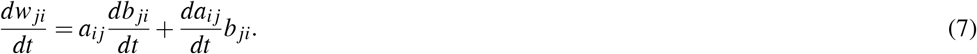

Here, we first consider the homeostatic plasticity of synaptic efficacy to maintain the overall stability of neuronal activity, referred to as synaptic scaling, in order to modify the synapse. It has been shown that neurons regulate their excitatory synaptic weights to achieve the biologically desired firing rate, *r*_0_^18, 32^. It allows a neuron *j* to decrease or increase synaptic effectiveness when its long-term firing rate, *r* _*j*_, is greater or less than *r*_0_.

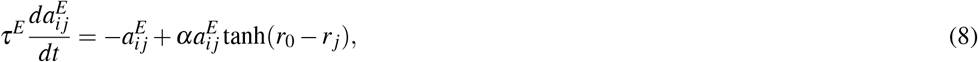

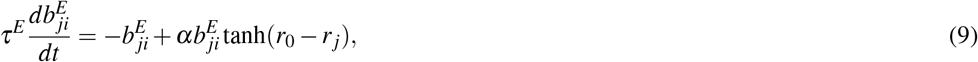

where *α* and *τ*_*E*_ are the scaling factor and time constant, respectively. The long-term firing rate *r* _*j*_ is determined by

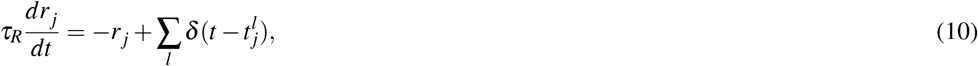

where *τ*_*R*_ is the time constant for the long-term firing rate and 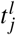 is the time a spike is elicited in output neuron *j*. While synaptic scaling is effective in stabilizing the firing activity of a network, it is insufficient for learning the information encoded in the input. Therefore, a correlation-based learning rule inspired by the N-Methyl-D-Aspartate (NMDA) receptor^11–13^ has been proposed in previous works^14, 15^. In the weakly supervised learning rule^14^, neurons are able to learn embedded patterns, as well as in^15^, a version following Dale’s rules, which operates in an unsupervised manner.

In more detail, the correlation between the input from pre-synaptic neuron *i* and output membrane potential of post-synaptic neuron *j*, denoted as *V*_*j*_(*t*), provides the signal for modifying the synaptic strength, denoted as *ε*_*i j*_(*t*) which is referred to as eligibility^14^:

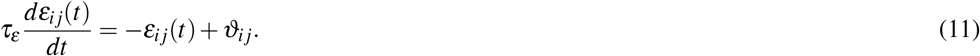

In this context, *ϑ*_*i j*_(*t*) represents the correlation between the input from the afferent *i* and the output neuron *j*, reflected by the membrane potential *V*_*j*_.

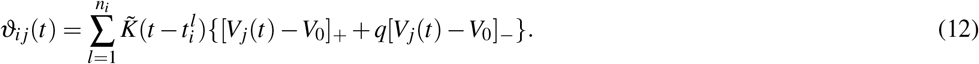

*V*_0_ represents the modification threshold^33^ and is consistently set to zero. The parameter *q* takes a value of 0 for excitatory afferents and a value of 1 for inhibitory afferents^15^. During each time step, all spikes *n*_*i*_ in the *i*-th afferent (elicited before time *t*) are employed to update the synapses. Also note that the kernel used to compute the correlation 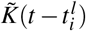 is given by the initial(input)kernel 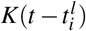 convolved with the kernel of the leaky integrate and fire equation 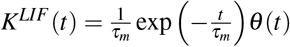, so the actual effect of an input spike on the post-synaptic membrane potential. The analytical expression 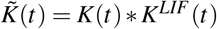 of this kernel is given by

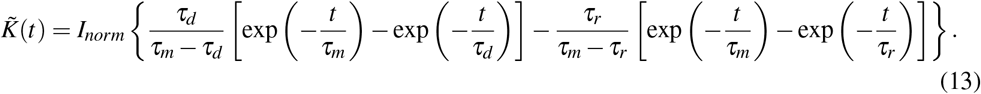

While the learning signal in excitatory synapses is positive, hetero-synaptic plasticity is used to provide negative changes, inducing selectivity and specificity. Here, we consider that the resources required to enhance the pre-and post-synaptic components *a*_*i j*_ and *b* _*ji*_ of the weights *w*_*ji*_ = *a*_*i j*_*b* _*ji*_ are inherently limited. As a result, we propose that these resources are distributed competitively, reflecting the limited availability and competitive nature of their allocation. The heterosynaptic plasticity of both pre- and post-synaptic regions is a function of the induced eligibility signal. The post-synaptic heterosynaptic plasticity, is implemented by subtracting the mean of the eligibilities. It affects changes in the post-synaptic components *b* _*ji*_:

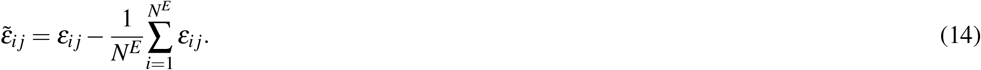

Nevertheless, it is hypothesized that the signals governing the changes in the pre-synaptic components *a*_*i j*_ depend on the magnitude of the potentiation observed on the post-synaptic side^34^. The signal for pre-synaptic competition is given by the signal for postsynaptic potentiation 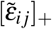, diminished by the mean of the potentiation signals of all other postsynaptic synapses, however, only in case more than one postsynaptic component receives a signal for strengthening. This is implemented by the following:

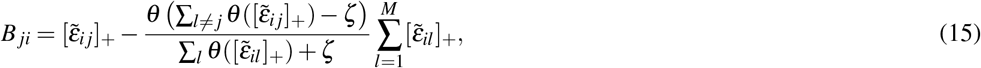

where *θ* is the Heaviside function, *ζ* is a small positive number (*ζ <<* 1), and M is the number of post-synaptic neurons. When applying our model to highly synchronized data, such as speech recordings, we attempted to increase synaptic competition so that a reduced number of synapses undergo significant growth. This was done by making the following modification:

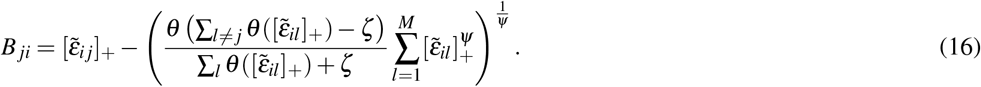

where *ψ* is the competition enhancement parameter. Increasing the parameter *ψ* beyond 1 enforces competition between synapses. In other words, a *ψ* value greater than one results in a greater degree of synaptic competition than when *ψ* is equal to one (Jensen Inequality). With this we model the dynamics of the synaptic components *a* and *b*:

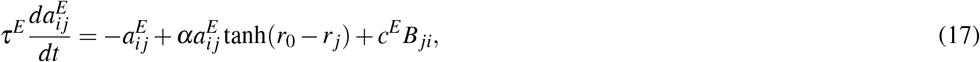

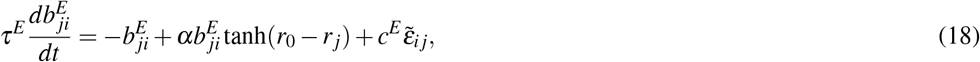

where *τ*_*E*_ is the time constant for changes of excitatory synapses and *c*^*E*^ the learning rate. For inhibitory neurons we use:

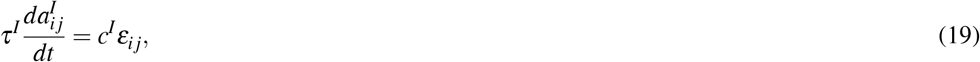

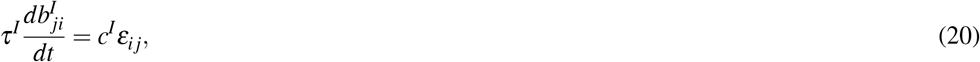

where *τ*_*I*_ is the time constant for inhibitory synapses and *c*^*I*^ is the learning rate. We consider *c*^*E*^ to be smaller than *c*^*I*^; therefore, inhibitory neurons can adapt more quickly to maintain balance at both global and detailed levels, and excitation cannot conflict with the relatively slow limiting mechanism of synaptic scaling^15^.

The phenomenon of simultaneous and faster Hebbian plasticity of inhibition is essential for maintaining a global balance within neural systems. Thereby the neuron is placed in a fluctuating regime in which the excitatory and inhibitory weights exhibit significant strength, resulting in substantial membrane potential fluctuations. Once this balance is achieved, further weight adjustments driven by the stochastic background restrict the weights to a fixed-point balance via a random walk process. Furthermore, the integration of a frequent pattern induces an additional drift that systematically shifts the weights away from the fixed point until the target number of spikes is induced by the pattern alone, independent of the response of the background activity ^15, 35^.

### Simulation Details

In order to reduce the computational effort, the network was trained epoch-wise. That means that after receiving input for a cycle of a given epoch length *T* ms, the changes in the synapses were calculated. Therefore, Eq. 12 was replaced by

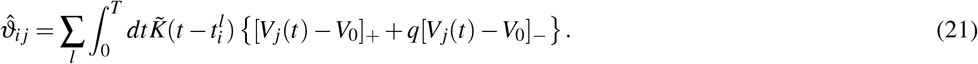

The low-pass filtering equations for the eligibility Eq.11 and the long-term firing rate Eq.10 were replaced by moving averages

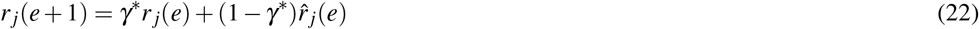

And

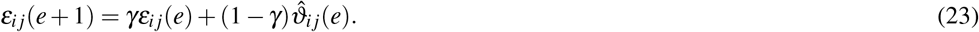

Here 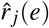 and 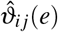 indicate the contributions to the long-term firing rate and eligibility of the current epoch *e* respectively with the initial conditions *r* _*j*_(0) = 0 and *ε*_*i j*_(0) = 0. *γ* and *γ*^*^ denote the filtering strength and are therefore parameters that depend on the respective timeconstants from Eq.10 and Eq.11. This parameter limits the learnability of patterns that are rarely presented^15^. All the parameters used in the simulations can be found in table 1.

**Table 1.** List of parameters.

| Symbol | Description | Value |
| --- | --- | --- |
| $\tau_m$ | Membrane time constant | 15 ms |
| $R_m$ | Membrane resistance | 1 |
| $\tau_r^E$ | Rise time of excitatory currents | 0.5 ms |
| $\tau_r^I$ | Rise time of inhibitory currents | 1 ms |
| $\tau_d^E$ | Decay time of excitatory currents | 3 ms |
| $\tau_d^I$ | Decay time of inhibitory currents | 5 ms |
| $r^E$ | Rate of excitatory neurons | 5 Hz |
| $r^I$ | Rate of inhibitory neurons | 20 Hz |
| $N$ | Number of pre-synaptic neurons | 500 |
| $N^E$ | Number of pre-synaptic excitatory neurons | 400 |
| $N^I$ | Number of pre-synaptic inhibitory neurons | 100 |
| $c^I$ | Inhibitory learning rate | $10^{-3}$ |
| $c^E$ | Excitatory learning rate | $0.9 \times 10^{-3}$ |
| $\gamma$ | Long term firing parameter | 0.9 |
| $\gamma^*$ | Eligibility parameter | 0.99 |
| $\Delta t$ | Time step | 0.1 ms |
| $\alpha$ | Scaling factor | 0.01 |
| $\alpha$ | Scaling factor (real data) | 0.04 |
| $T$ | Epoch duration (single neuron) | 2000 ms |
| $T$ | Epoch duration (Neural Network) | 1000 ms |
| $T$ | Epoch duration (real data) | 2500 ms |
| $\chi$ | Decay term | $10^{-4}$ |
| $\psi$ | competition enhancement parameter | 12 |

**Table 2.**
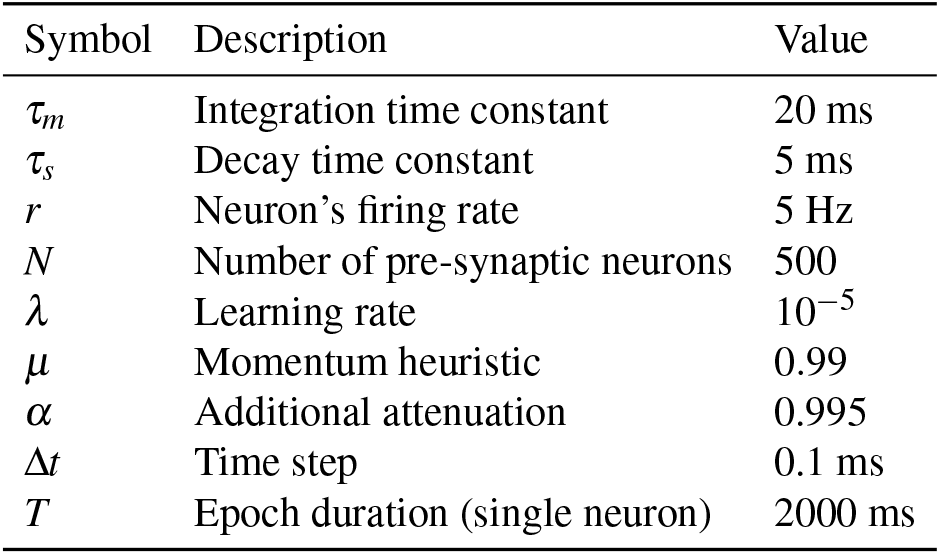
Parameters used for multispike-tempotron & correlation-based learning.

Due to structural limitations and other factors, synaptic strength cannot increase indefinitely. As a result, we restrict weight changes to prevent excitatory synapses from exceeding the upper limit of one.

Dale’s rule states that excitatory and inhibitory synapses cannot convert into each other. Therefore, we consider synaptic weights to be zero if weight changes during learning would change their type. This ensures that subtracting the mean in Eq.14 and Eq.15 does not affect the type of a synapse. We use the Euler method with a time step of Δ*t* for the numerical integration of Eq.13. Note, for the network model, the initial synaptic efficacies *a*_*i j*_ and *b* _*ji*_ are drawn from a Gaussian distribution with a mean of 0.1 and 10^−2^ standard deviations, while negative values are set to zero. In the single post-synaptic neuron model, they are drawn from a Gaussian distribution with a mean of 10^−2^ and standard deviations of 10^−3^, with negative values set to zero. In cases where there is only a single post-synaptic neuron, all values of *a*_*i j*_ are initially set to 1. Since there is no pre-synaptic hetero-synaptic plasticity, these *a*_*i j*_ values will remain at their initial value of 1, regardless of the number of learning cycles.

In the epoch-based approach, we use the following equations, which can be linearized to obtain the continuous version:

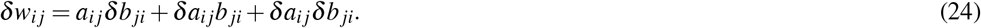

#### Inhibitory synapses

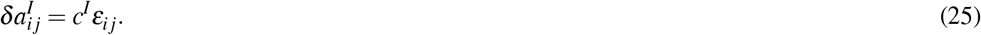

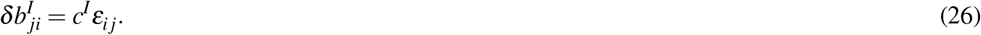

#### Excitatory synapses

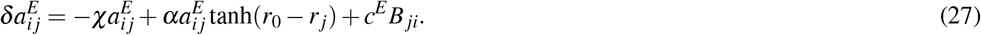

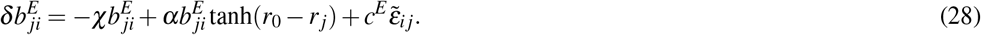

The additional factor *χ* in front of the decay term is given by the excitatory time constant *τ*_*E*_ . The other parameters *α* and *c*^*E*^ have been updated by that same factor as well.

### Noise robustness

#### Generation of noise

Noise was introduced in the form of Poisson spike rates. These were generated by computing the convolution of the initial (frozen) spike train with a Gaussian, where the standard deviation *σ* of the Gaussian represents the temporal jitter. This convolution yields a firing rate which then is used to generate the new spike train by another Poisson process . This process is visualised in Figure 11.

**Figure 11.**
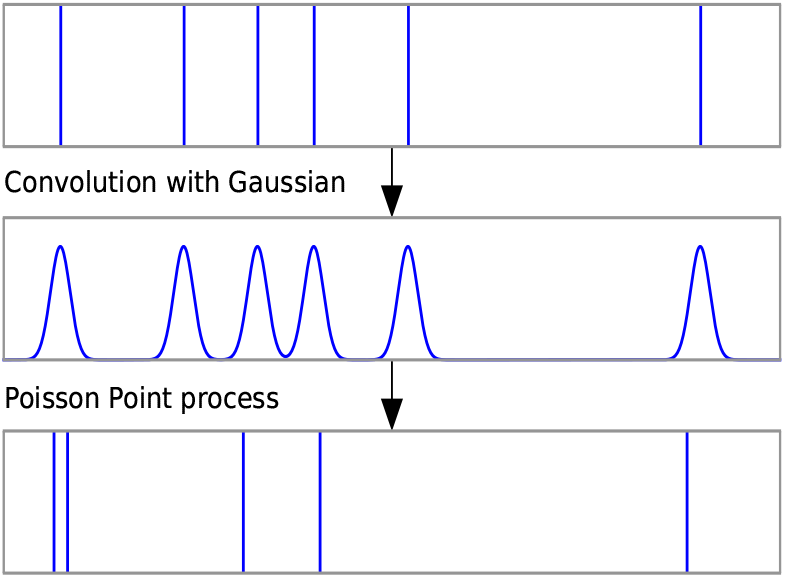
Schematic procedure of generation of the Poisson spike rates. Top: Initial spike train. Center: Convolution with the Gaussian. Bottom: New spike train drawn from the distribution of the center graph in a Poisson Point process.

#### Comparison to other models

When comparing our model to the approach of the multispike-tempotron and correlation-based learning the parameters listed in Tab. 2 were used. In addition to the learning rules of the respective algorithms, there was also a momentum heuristic and an additional attenuation implemented as in^14^.

## Code Availability

The source code supporting the findings of this study is available in the GitHub repository. The repository can be accessed at the following URL: https://github.com/MohammadDehghaniH/Incremental_Self-Organization_of_Spatio-Temporal_Spike_Pattern_Detection

## Author contributions

M.D-H. and K.P. designed research. M.D-H., L.M., and K.P. performed research. M.D-H., L.M., and K.P. wrote the paper.

## Acknowledgements

We would like to thank David Rotermund for his assistance in providing the spiked audio data. We acknowledge support by the following grants: DFG - grant (PA 569/5-1 to KP) During writing we took advantage from special tools including Bing Chat: https://www.bing.com, DeepL: https://www.deepl.com and ChatGPT.

